# A Ratio-Dependent Model of Replicator-Genetic Parasite Coevolution Demonstrates Instability of the Parasite-Free State

**DOI:** 10.1101/2021.02.20.432109

**Authors:** Faina Berezovskaya, Georgy P. Karev, Eugene V. Koonin

## Abstract

Nearly all organisms on earth are hosts to diverse genetic parasites including viruses and various types of mobile genetic elements. The emergence and persistence of genetic parasites was hypothesized to be an intrinsic feature of biological evolution. Here we examine this proposition by analysis of a ratio-dependent Lotka-Volterra type model of replicator(host)-parasite coevolution where the evolutionary outcome depends on the ratio of the host and parasite numbers. In a large, unbounded domain of the space of the model parameters, which include the replicator carrying capacity, the damage inflicted by the parasite, the replicative advantage of the parasites, and its mortality rate, the parasite-free equilibrium takes the form of a saddle and accordingly is unstable. Therefore, the evolutionary outcome is either the stable coexistence of the replicator and the parasite or extinction of both. Thus, the results of ratio-dependent model analysis are compatible with the hypothesis that genetic parasites are inherent to life.

## Introduction

With few exceptions, all organisms are afflicted by genetic parasites including viruses, transposons and plasmids [1–3]. The emergence of genetic parasites appears to be an inherent feature of replicator systems. Indeed, as soon as, in the case of a simplest possible replicator, such as a small RNA genome, there is an alienable resource required for replication, such as a replicase, parasites will emerge that will cheat by avoiding producing that resource and instead appropriating it from those replicators that do [4]. In this framework, the parasite is represented by a defective variant of the replicator. The emergence of such minimal genetic parasites is not a purely theoretical construct but closely recapitulates the classic experiments of Spiegelman and colleagues on *in vitro* evolution of RNA bacteriophage genomes as well as the evolution of defective interfering variants of numerous viruses [5–7].

Analysis of mathematical models of the evolutionary dynamics in predator-prey or host-parasite systems has been an important area in theoretical ecology ever since the classical Lotka-Volterra equations have been developed [8]. The crucial element of these models is the “trophic function” *g*(*R*, *P*) that describes the number of preys consumed per predator per unit time depending on the number of prey *R* and predators *P*. A variety of different trophic functions have been considered [9, 10].

Many early studies have focused solely on the dependency of the trophic function on the prey density, ignoring the effect of predator density; such models are known as “prey-dependent”. However, it has been recognized that the predator density could have a direct effect on the trophic function as well, and several predator-dependent models have been proposed [11]. The classical prey-dependent predator–prey models exhibit the so called “paradox of enrichment” [12–14], whereby increased resources cause a collapse of an ecosystem, and “biological control paradox” [15, 16] where efficient control agents cause severe pest outbreaks. These paradoxical phenomena are not captured by the prey-dependent or predator-dependent models, due to the simplifying assumptions on trophic functions, suggesting that more realistic coevolution models have to be developed.

Getz, Arditi and Ginzburg, and Berryman independently proposed ratio-dependent predator-prey models [17, 18] (reviewed in [9]), where the trophic function is set to depend on the single variable, the ratio *R*/*P*, rather than on the two separate variables *R* and *P*, i.e. *g*(*R*, *P*) = *g*(*R*/*P*). Using a standard incidence function, one gets

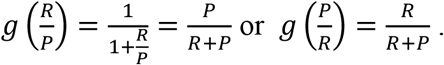

In the ratio-dependent models, the trophic function, that is, the interaction between prey and predators or between replicators and parasites, is represented by the term 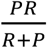, in contrast to the Volterra-type model where it is represented by the term *PR*. To emphasize the difference, let *R*~*const*, *P* → ∞; then, ~*P*, and 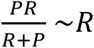; let *P*~*const*, *R* → ∞ then, *RP* ~*R*, and 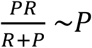. Accordingly, the two models show fundamentally different dynamics when the numbers of replicators and parasites are of different orders.

Importantly, the ratio-dependent models do not produce the paradox of enrichment and biological control paradox (for details, see [9, 19]).

Although there have been considerable debates over the relative strengths and weaknesses of prey-dependent and ratio-dependent models [20–22], the latter have enabled mathematical description of some key experimental observations [9, 10, 19, 23–25]. In particular, ratio-dependent models have been instrumental in explaining such phenomena as the extinction or coexistence of the prey or predator populations depending on the ratio of the initial population sizes (see, e.g., [26, 27]. The classical prey-dependent models are not fit to address these effects. These models fail to accurately describe the outcomes of biological control efforts where an attempt is made to control the population of a pest by introducing a parasite (a control agent). Under this biological control paradox, highly efficient control agents cause extreme pest outbreaks because such agents overexploit the resource and die out, allowing the pest population to rapidly recover [15, 16]. This behavior, which is essential to consider for planning biological control, can only be captured by ratio-dependent models.

The standard Lotka-Volterra trophical function works well when the abundances of prey and predator are of the same order. The ratio-dependent version is far more appropriate when abundances of the prey and predator (or host and parasite) are of different orders, for example, many predators and not enough prey. This is the case for the simplest models of replicator - genetic parasite systems [4] because, at the time of the parasite emergence, the parasite is bound to be far less numerous than the host replicators. Considering this feature of the system could be critical to obtain an accurate description of the evolutionary dynamics of replicator - genetic parasite systems.

The ratio-dependent models present many interesting mathematical features [28–34]. A complete parametric analysis of the stability conditions and dynamics of a ratio-dependent model with logistic growth of prey has been reported [25] as well as a model with the Allee-type growth of prey [35]. The principal distinguishing property of such models is that the origin is a singular equilibrium point whose characteristics (depending on parameters) crucially determine the behavior of the model. Ratio-dependent models can have complicated, rich dynamics and display distinct dynamic properties that have never been observed in simple two-dimensional predator-prey models. For example, the origin can be simultaneously attractive and repulsive in different sectors of its neighborhood, thus shedding new light on the ways of the system extinction. In particular, the models can demonstrate the regime of “deterministic extinction” [25, 29, 31]. Coexistence of several dynamic regimes with the same set of parameters depending on the starting point in an arbitrary small vicinity of origin can also be observed.

The goal of this work is to perform the mathematical analysis of a ratio-dependent version of the models of replicator–genetic parasites system [4] and its Volterra-type modifications [36]. Analyzing the ratio-dependent version of the model, we show here that the system can exist in a stable non-trivial state with non-zero numbers of both replicators and genetic parasites in a large domain of the model parameters. These findings support the hypothesis that emergence and persistence of genetic parasites is an intrinsic feature of evolving replicators that is caused by the evolutionary instability of parasite-free states [4].

### 1. The models

The main assumptions of the previously proposed conceptual model [4] are as follows. The system consists of two populations: autonomous replicators (*R*), and genetic parasites (*P*) that lack autonomous replication capacity. The kinetics of replication depends on the interaction between the template (either the host replicator or a parasite) and the replicase, which is produced by the replicator but not by the parasite. Thus, the replication rate is proportional to the product of the respective population sizes, that is, *R*^2^ for the replicator and *RP* for the parasite. The decay rates are constant (*e*_*R*_ and *e*_*P*_) for the replicator and the parasite, respectively. The parasite replicates faster than the replicator by the factor *q* ≥ 1 which can be interpreted as the “parasite advantage” parameter (in the simplest possible terms, the parasite genome is *q* times shorter than the replicator, so its replication is faster by the factor of *q*). Both populations are environmentally limited by the same resources; the parasites consume *q* times less resources per individual compared to the host replicator; the environment carrying capacity *K* determines the point where replication becomes resource-limited. The replicators possess a (costly) defense mechanism with efficiency *e* ≥ 0 that can suppress the parasite replication by a factor 1 + *e*, at the cost to the replicator’s replication rate 1 + *αe* (where *α* ≥ 0 is the cost factor of defense factor).

The dynamics of such a system is described by the following pair of ordinary differential equations:

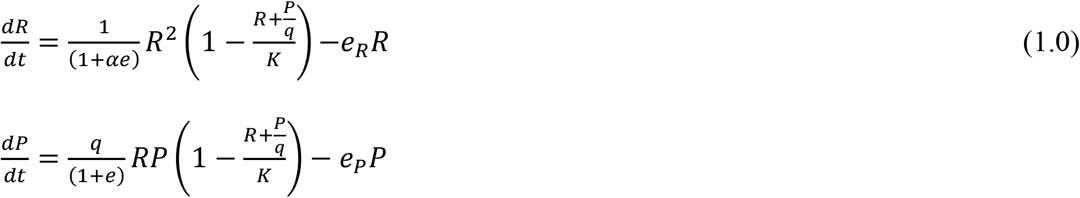

where *K* is the carrying capacity of the replicator population, and *Kq* is the carrying capacity of the parasite population. However, subsequent analysis has shown that model (1.0) has non-trivial equilibria only for special values of parameters in which case a line of non-isolated equilibria appears.

A Volterra-type modification of model (1.0) has been proposed and studied [36]:

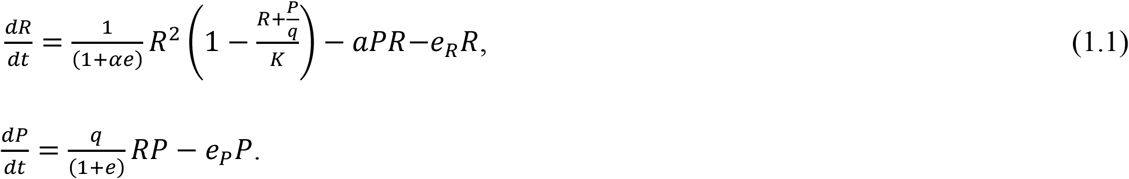

In this model, the “ecology” of the system changes: the number of replicators (not a common resource) is the only limiting factor for the reproduction of the parasites. It is further assumed that parasites can damage the replicator population; the additional mortality rate is given by the term *αPR*. It was proved that the (*R*, *P*)-system (1.1) can exist in a stable state, either stationary or oscillation, within a bounded area of parameters [36]. There also exist regimes where either the population of the parasite or the system as a whole goes extinct. Thus, under this model, the emergence and persistence of genetic parasites is possible but not inevitable.

Here we propose and study a modification of model (1.1) based on <u>r</u>atio-<u>d</u>ependent (RD) interaction between replicators and parasites. The model is of the form

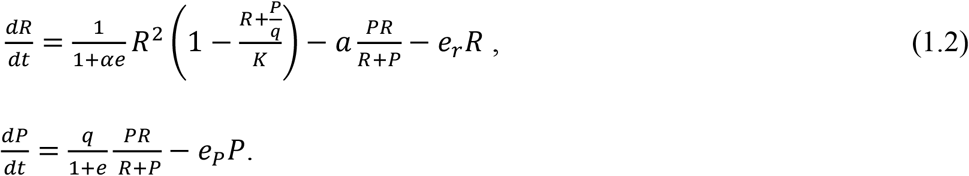

Using the ratio-dependent version (1.2) of the replicator-genetic parasites model, we show that the system can exist in a stable equilibrium with non-zero numbers of both replicators and genetic parasites in a very large domain of the model parameters. This result supports the hypothesis of inevitable emergence and persistence of genetic parasites caused by the evolutionary instability of parasite-free states [4]. We study also the role of parameters in the existence and collapse of the system.

The system (1.2) has a singular point *O*(*R* = *P* = 0). To avoid this singularity, let us change the time:

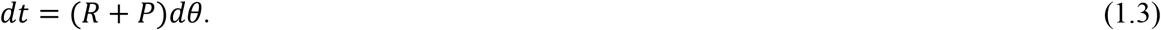

This transformation changes the velocity of the system’s movement along trajectories according to the new time *θ* but does not change their shapes. Then, we obtain the system

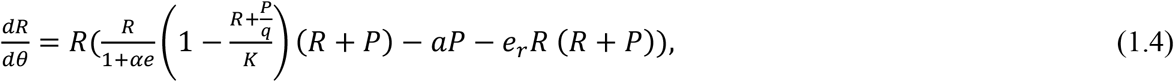

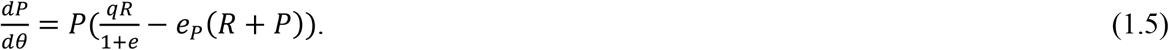

Denoting

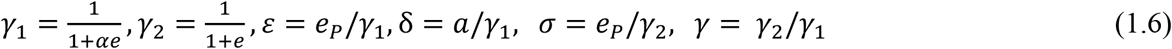

and changing again the independent variable

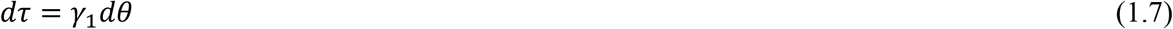

we can write the system in a more convenient form

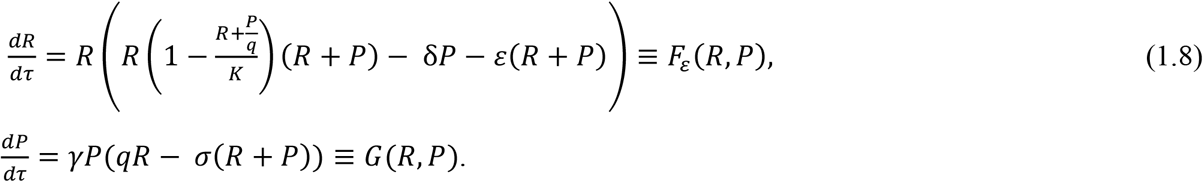

where 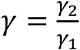.

The model (1.8) is structurally stable in the positive quadrant (*R*, *P*) according to the following proposition.

#### Proposition 1.

*The phase curve* {*R*(*t*), *P*(*t*)} *of model (1.8) stays in the bounded domain of positive quadrant* (*R*, *P*) *indefinitely if the initial values R*(*t* = 0) ≥ 0*, P*(*t* = 0) ≥ 0.

See Appendix 1 for proof.

In what follows, we investigate phase-parameter dynamics of the system (1.8). We firstly study the model (1.8) with the parameter *ε* = 0 (see system (2.1) below), and then, analyze the role of parameter *ε* in the model dynamics.

### 2. Analysis of the model with *ε* = 0. Equilibrium states

Consider the system

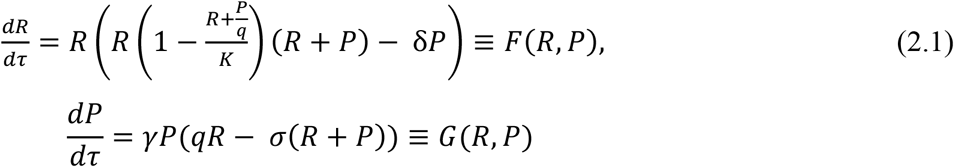

which coincides with the model (1.8) if *ε* = 0.

Null-clines of model (2.1) are given by the system

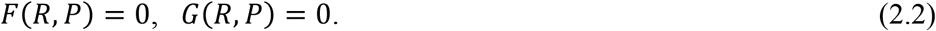

The system (2.1) has trivial null-clines *P* = 0, *R* = 0 and non-trivial positive null-clines

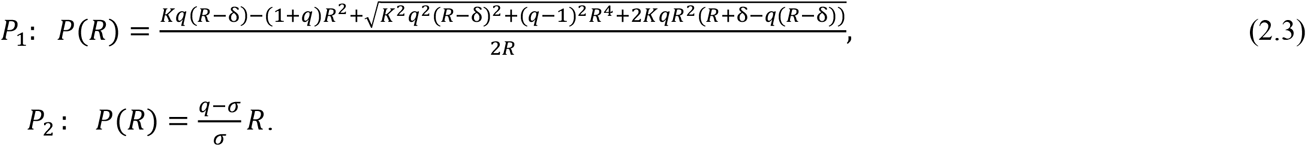

One can show (expanding the right part of equation (2.3) in a series with respect to *R*) that the isocline *P*_1_ touches the *R* axis in the point *O* because *P*_1_ (*R* = 0) = 0 and 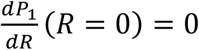 and intersects the axis in the point *C*(*K*, 0), with the tangent 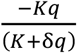 (Fig. 1).

**Fig 1.**
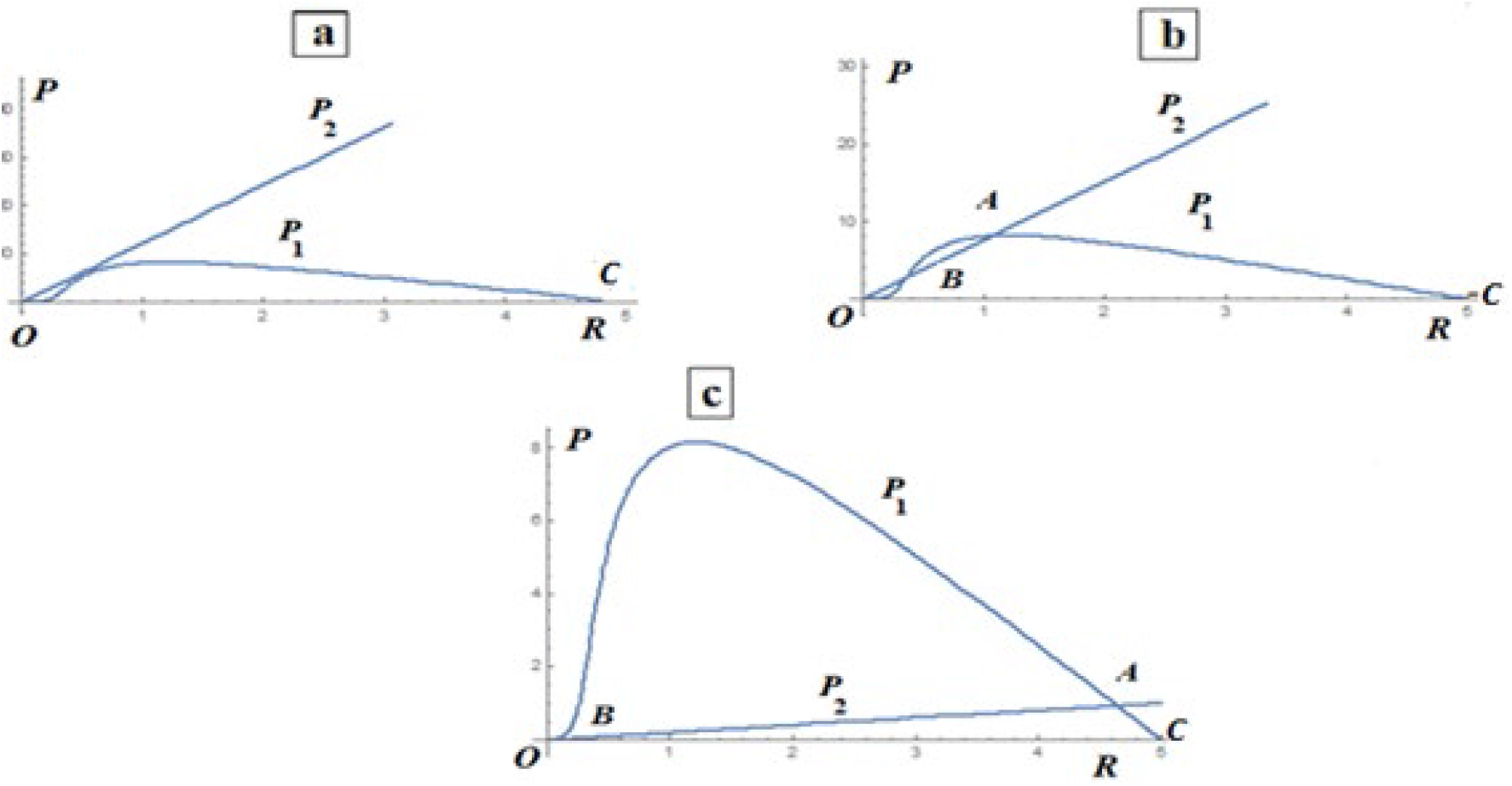
Nontrivial null-clines *P*_1_(*R*), *P*_2_(*R*) of model (2.1) given by formulas (2.3) where *K* = 5, *q* = 3, δ = .3, γ=1; **a:***σ* = .23; **b:***σ* = .35, non − trivial equilibria *A*(2.06, 8.07), *B*(. 35, 2.67); **c**: *σ* = 2.5, equilibria *A*(4.64, .93), *B*(. 05, .01).

The points of intersection of null-clines are the equilibria of the model. The model can have up to four equilibria: *O*(0,0), *C*(*K*, 0), *A*(*R*_A_,*P*_A_), *B*(*R*_B_,*P*_B_) (Fig. 1) where

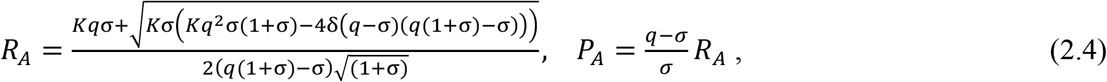

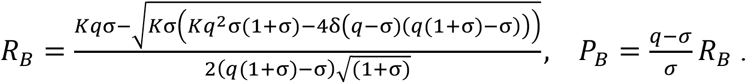

It can be verified by simple algebra that positive, non-trivial equilibria *A, B* exist if the following inequalities hold

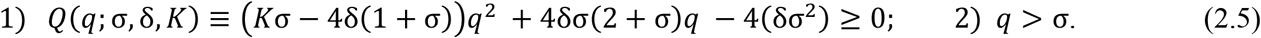

The first inequality (2.5) can be conveniently expressed through the roots *q*_1_, *q*_2_ of the polynomial *Q*(*q*; σ, δ, *K*), which can be presented in the form *Q*(*q*; σ, δ, *K*) ≡ (*K*σ − 4δ(1 + σ))(*q* − *q*_1_)(*q* − *q*_2_)

where

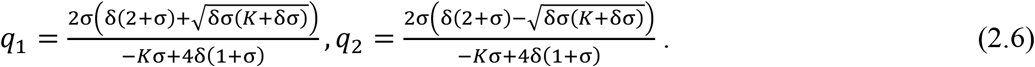

Note that *q*_2_ < *q*_1_ if (*K*σ − 4δ(1 + σ)) < 0 and *q*_2_ > 0 > *q*_1_ if (*K*σ − 4δ(1 + σ)) > 0.

Then, for positive *q, Q*(*q*; σ, δ, *K*) ≥ 0

if (*K*σ − 4δ(1 + σ)) < 0 and *q*_2_ < *q* < *q*_1_ or if (*K*σ − 4δ(1 + σ)) > 0 and *q* > *q*_2_.

One can easily see that σ > *q*_2_. Then, both inequalities (2.5) hold if

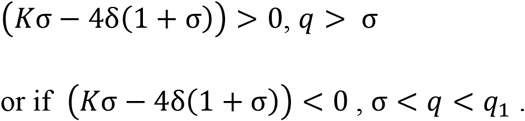

For any fixed value *K*, the line

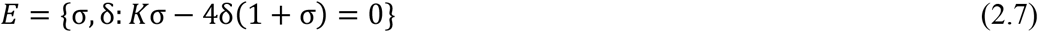

divides the parameter space (σ, δ) into two domains (see Fig.2A).

**Figure 2A.**
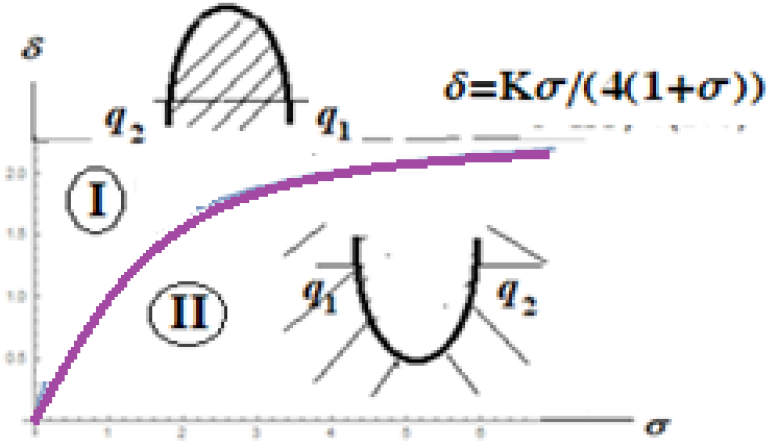
{δ, σ} −cut of the parametric portrait, the line ***E***: *K*σ − 4δ(1 + σ) = 0 (violet).

The corresponding domains in {*σ*, *q*} −cut of the parametric portrait are shown in Figure 2B:

**Figure 2B.**
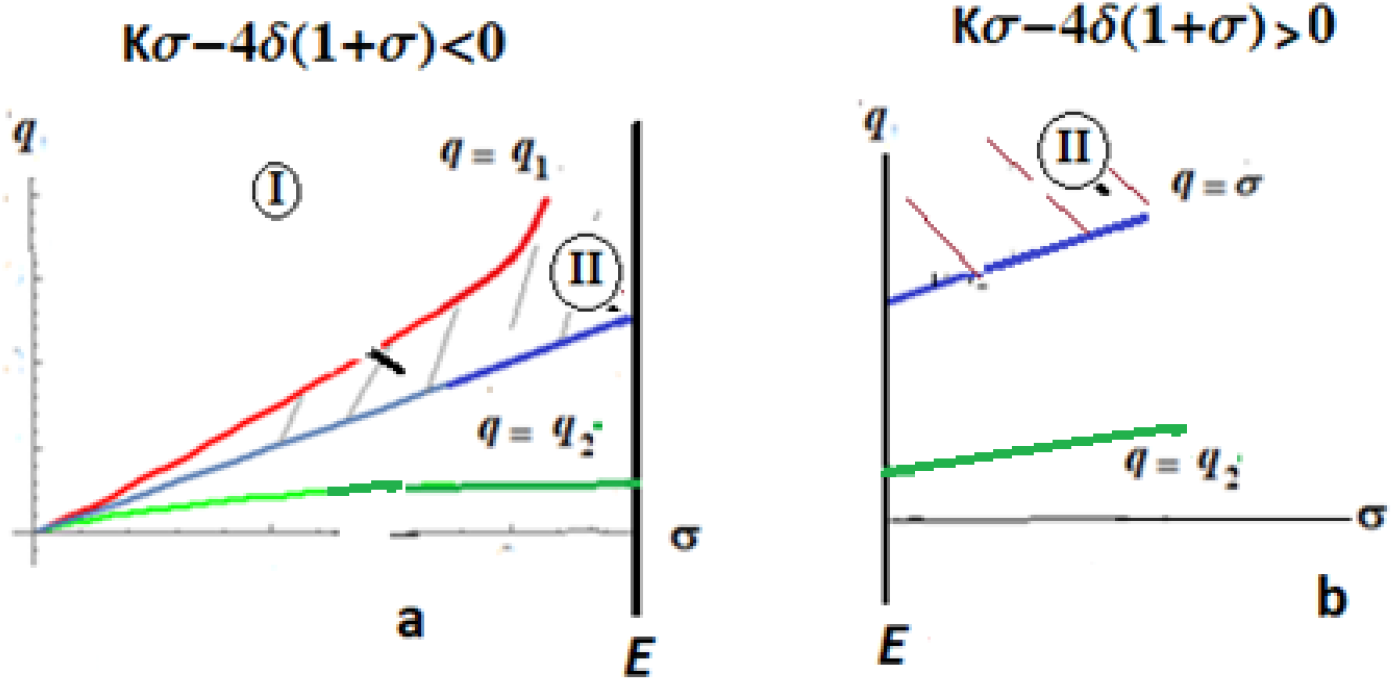
{σ, *q*} −cut of the parametric portrait for (**a**) *Kσ* − 4*δ*(1 + *σ*) < 0, (**b**) *Kσ* − 4*δ*(1 + *σ*) > 0. The line 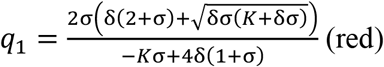, the line 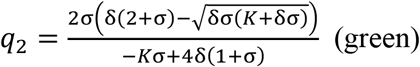. Domains I and II contain 2 and 4 non-negative equilibria, respectively. The system has positive equilibria *A, B* in domain I for σ < *q* ≤ *q*_1_ and in domain II for σ < *q*; the areas of existence of *A, B* are shadowed.

The following Proposition gives conditions for the existence of equilibria of the evolving system.

#### Proposition 2

(see Fig.1).

*In the first quadrant of plane* (*R*, *P*) *system (2.1) has equilibria*

1. *O*(0,0), *C*(*K*, 0) *for all parameter values, and*
2. *A*(*R*_*A*_, *P*_*A*_), *B*(*R*_*B*_, *P*_*B*_) *with coordinates (2.4) for (a) any* 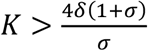 *and q > σ*; (*b*) 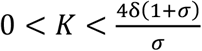 *and σ* < *q* < *q*_1_ *where q*_1_ *is given by* (2.6) (see Figs 2A, B);
3. *Equilibria A, B appear/disappear in the positive quadrant on the boundary **Q**: q* = *σ where the point A coincides with C and the point B coincides with O* (see Fig.2B).
4. *Positive equilibria A, B appear/merge composing unique point AB (R*_*AB*_, *P*_*AB*_) *with coordinates* 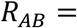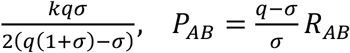 *in the boundary*

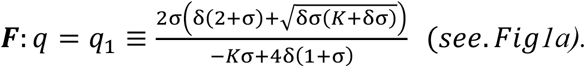

See Appendix 2 for proofs.

*Corollary.* The model has 4 non-negative equilibria only for *q* > σ in domain (II) (see Fig. 2). Here (I) is the domain where *K*σ − 4δ(1 + σ) > 0 (Fig. 2b), and (II) is the domain where σ − 4δ(1 + σ) < 0, *q* < *q*_1_ (Fig. 2a). These domains are separated by the boundary ***E***: *K*σ − 4δ(1 + σ) = 0.

The following statements describe the structures of equilibria (see Appendix 3 for proofs).

#### Proposition 3.

1. *The equilibrium C is a stable node for q* < *σ and a saddle for q* > *σ. For q* = *σ equilibrium C is a saddle-node with an attracting node sector for positive P. Crossing the line q* = *σ corresponds to the transcritical bifurcation in the system when the points C and A exchange of their stability (see, e.g., [37]*.
2. *The equilibrium B is a saddle, and the equilibrium A is a topological node (stable/unstable node/spiral or a center) for parameter values given in Proposition 2: *q* > σ if* 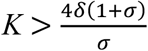 and *σ < q < q*_1_ *if* 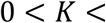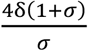
3. *If* 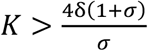 *then the equilibrium A is a stable node. If* 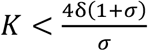 *then the equilibrium A can be a stable/unstable node or a stable/unstable spiral. The equilibrium A changes stability via supercritical Andronov-Hopf bifurcation that happens on the line*

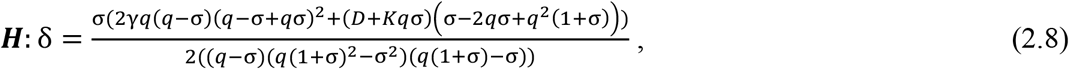

where 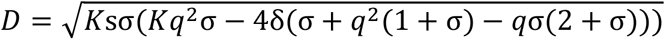
4. *Positive equilibria A, B merge composing unique point AB (R*_*AB*_, *P*_*AB*_) *with coordinates* 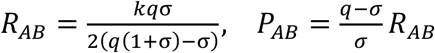 *in fold bifurcation that happens in the line*

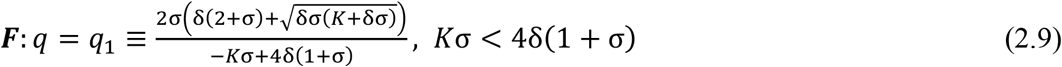

*Remark.* The system of equations (2.8), (2.9) defines implicitly the curves in {*σ*, *q*} – space when all other parameters are fixed; the curves form boundaries of the parametric portrait of system (2.1).

#### Proposition 4.

*Let q* > σ. *Then the positive vicinity of the origin O(0,0) of system* (2.1) *contains an elliptic sector* (see Fig. 3).

**Fig. 3.**
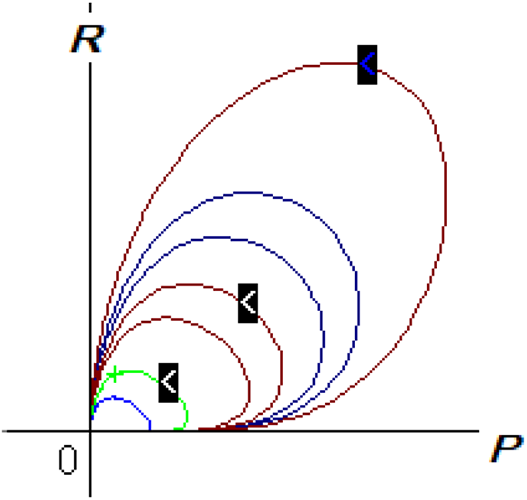
Structure of the equilibrium *O*(0,0) of system (2.1) for *K* = 10, *γ* = 1, *q* = 5.45, *σ* = 2, *δ* = 3; the vicinity of *O*(0,0) contains an elliptic sector (together with stable and unstable node sectors).

#### Proposition 5.

*In the parametric domain* 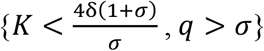, *there exist values of the system parameters such that*

1. *outcoming separatrix of the saddle C coincides with incoming separatrix of the saddle B* (see Fig.1b). *This non-local bifurcation occurs on the line **L**(see Figures 4,5)*;
2. *incoming and outcoming separatrices of the saddle B form a separatrix loop containing point A inside*. *This non-local bifurcation occurs on the line PS (see Figures 4,5).*

Statements of Proposition 5 were verified numerically.

We summarize our results in the following theorem.

**Fig. 4.**
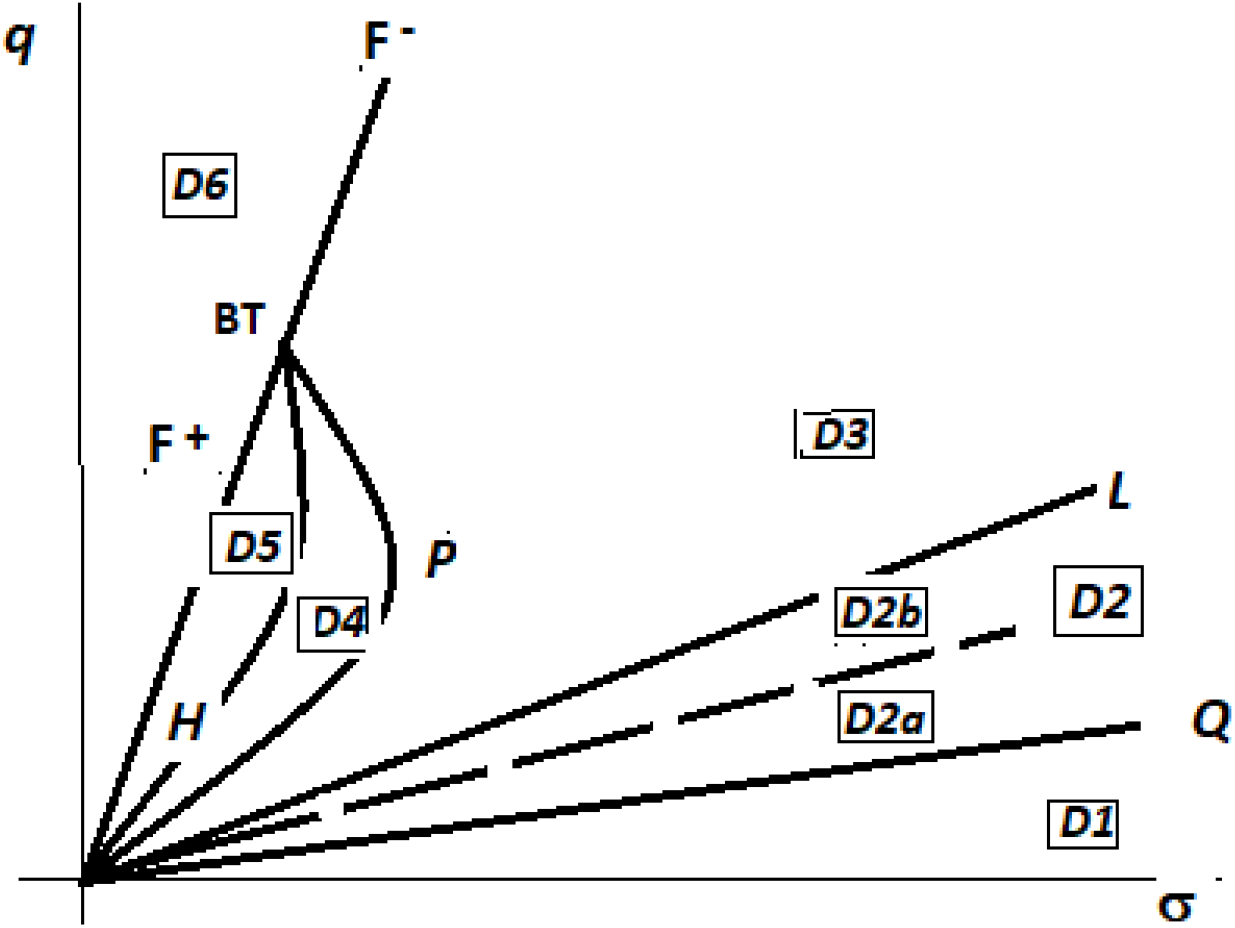
Schematic parameter portrait of model (2.1) representing by {*σ*, *q*} − cut of its {*K*, *δ*, *σ*, *q*, *γ*} − parameter portrait for any fixed *K* > 0 and “typical” values δ, *γ*.

**Fig. 5.**
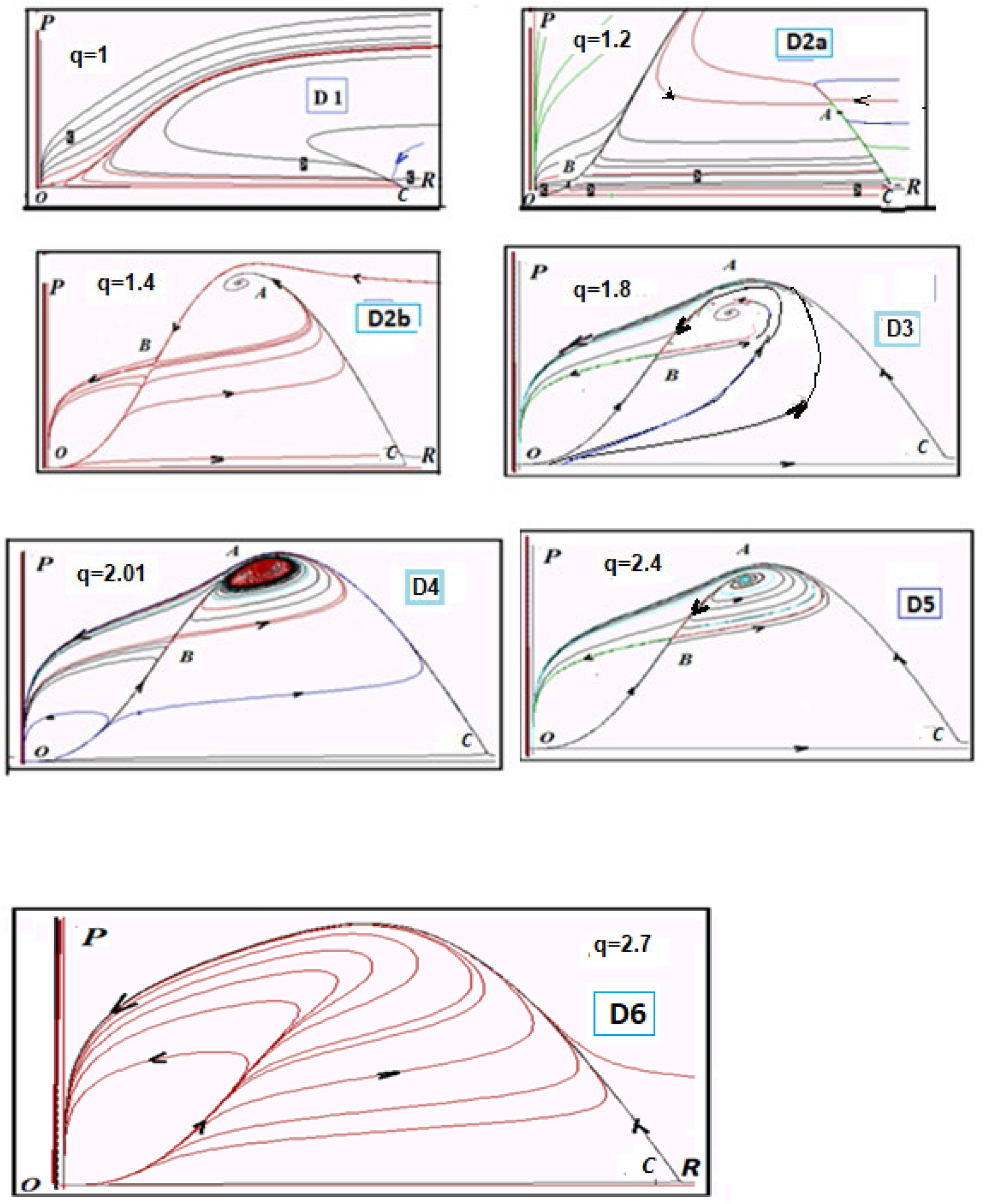
Phase portraits of model (2.1) in domains shown in Fig. 4 as *K*=10, {*δ*, *σ*} = {3, 1}, *γ* = 1. The values of the parameter *q* are shown on the Figures.

#### Theorem 1.

1. *For any non-negative parameters in the non-negative quadrant of* {***R***, ***P***} −*plane System (2.1) can have from two*, O(0,0), C(***K*** *,0,*), *up to four equilibria, O, C, A, B. Equilibria A and B exist in the positive quadrant of* {***R***, ***P***} − *plane if* ***q*** > ***σ****. Coordinates of the points A, B are given by formulas (2.4). The point C is a saddle if* ***q*** < ***σ*** *and a stable node if* ***q*** > ***σ***; *the point B is a saddle; the point A is topological node/ spiral, point O is non-hyperbolic for all parameter values.*
2. *Parametric and phase portraits of system (2.1) are given in Fig-s. 4 and 5 for any fixed parameter K>0 and non-negative values of parameters* {*δ*, *σ*, *q*, *γ*}. *Figure 4 schematically presents* {*σ*, *q*} − *cut of* {*K*, *δ*, *σ*, *q*, *γ*} − *parameter portrait of the model for some fixed values of K*, *δ*, *γ; the boundaries of domains correspond to the lines where bifurcations of co-dimension 1 happen in the system. Figure 5 shows the* {*R*, *P*} −*phase portraits of the system that realize in the respective domains of the parameter apace are shown in Fig. 4.*

Let us consider in detail the parametric portrait in Fig. 4.

The {*K*, *δ*, *σ*, *q*, *γ*} − boundaries of the parametric portrait are as follows.

1) ***Q***: *q* = *σ*.

The equilibria *A*, *B* appear in the first quadrant on the boundary *Q* where the point *A* coincides with *C* and the point *B* coincides with *O* (see Proposition 3).

2) ***E***:*Kσ* − 4δ(1+*σ*) = 0.

If *Kσ* − 4*δ*(1 + *σ*) > 0 and *q* > *σ*, then, the system has a positive equilibrium *A* that is a stable node.

If *Kσ* − 4*δ*(1 + *σ*) < 0, then, the system has the positive equilibrium *A* if *σ* < *q* < *q*_1_ (see Eq. 2.6), which can be stable or unstable node or spiral.

3) The fold boundary ***F = F ^−^*** ∪ ***F ^+^*** is given by the equation (2.9)

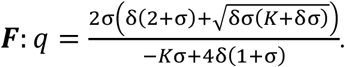

The boundary corresponds to the appearance /disappearance of non-trivial equilibria *A, B* in the positive quadrant of {*R*, *P*} − plane. On the boundary ***F^−^**,* the stable node *A* and the saddle *B* merge, producing a saddle-node *AB* with an attracting node sector. On the boundary ***F^+^**,* the unstable node *A* and the saddle *B* merge, producing a saddle-node *AB* with a repelling node sector. The point *BT* connecting ***F ^−^*** and ***F ^+^*** is a point of intersection of the lines ***F*** and ***H***; it corresponds to the Bogdanov-Takens bifurcation of co-dimension 2 where the equilibrium *AB* is a degenerated saddle in the phase plane (see, e.g., [38]). Coordinates of the point *BT* in parameter plane (*q*, σ) are the roots of the system {*F, H*}*=*0 given by formulas (2.8), (2.9) with *D*=0.

4) ***PS:*** the boundary (no analytical expression) corresponds to the non-local bifurcation ‘separatrix cycle’ which is composed of two separatrices of the saddle *B*; as *q* increases and intersects ***P*** (coming to D3), the separatrix cycle turns into an unstable cycle that contains the stable equilibrium *A* inside.

5) ***H***: the supercritical Andronov-Hopf bifurcation (changing stability of the point *A*) occurs on the boundary ***H*** (given by formulas (2.8)). On this boundary, an unstable limit cycle around the stable equilibrium *A* “shrinks” to the point *A* that becomes an unstable equilibrium.

6) ***L:*** this boundary (no analytical expression) corresponds to a non-local bifurcation of coincidence of the separatrices of the saddles *C* and *B.* Before this bifurcation, the outcoming separatrix of *C* enters to the point *A*, and after the bifurcation, it enters to the point *O.*

Note that Figures 4 and 5 demonstrate two different sequences, *I* and *II*, of qualitative behaviors of the system, with a switch occurring with the increase of the parameter *q* value at different fixed values of *σ*. The first sequence consists of portraits realized in the domains {D1, D2, D5, D6} and the second one consists of portraits in the domains {D1, D2, D3, D4, D6}.

In Domain *D1, q* < *σ*, the system has only two equilibria in the first quadrant of {*R*, *P*} − plane, namely, the saddle-node *O*, and the stable node *C*. The system also has only these two equilibria in domain *D6,* where {*q* > *q*_1_} ∪ { *Kσ* − 4*δ*(1 + *σ*)} < 0 and *q*_1_ is given by equation (2.10). However, in domain *D6,* the point *O* has an elliptic sector in its positive neighborhood, whereas the point *C* is a saddle (see Fig.5).

Domain *D2* is common for both sequences. In this domain as well as in domains *D3, D4* and *D5,* equilibrium *O* has an elliptic sector bounded by separatrices of the saddle *B.* In domain *D2* as well as in domains *D3, D4* there exists *non-trivial positive attractive point A*. Let us emphasize that with increasing of the value of parameter *q*, the area of attraction of the equilibria *O* increases whereas the area of attraction of the equilibria A decreases.

### 3. Model (1.8)

This model with *ε* > 0

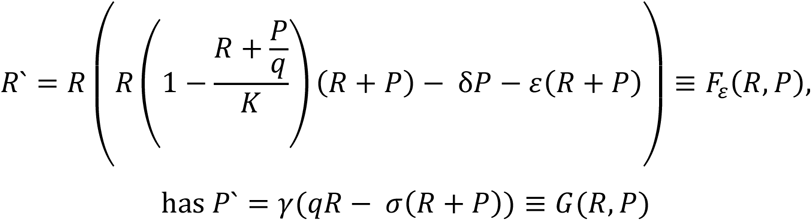

equilibria whose coordinates satisfy the system {*F*_*ε*_(*R*, *P*), *G*(*R*, *P*)} = 0.

The system has trivial null-clines *P* = 0, *R* = 0 and non-trivial positive null-clines

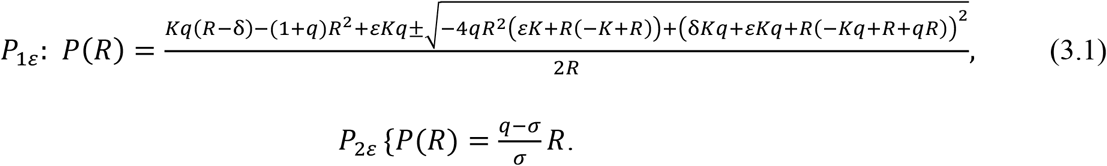

It is easy to see that, for small enough *ε*, the intersections of null-clines are similar to those in Fig. 1 where *ε* = 0 everywhere except the axis *R*. We showed that system (2.1) has no more than 4 equilibria, *A,B,C,O.* The system (1.8), however, can have 5 non-negative equilibria:

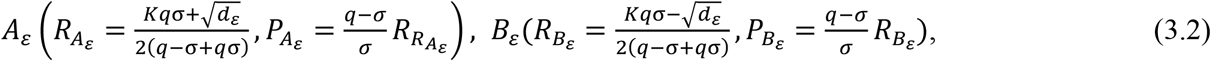

where *d_ε_* = *K*σ(*Kq^2^*σ − 4(*εq* + δ(*q* − σ))(*q* − σ + *q*σ)), and

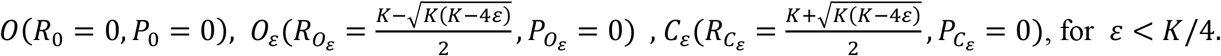

Expanding *R*-coordinates of the equilibrium points to series, we get

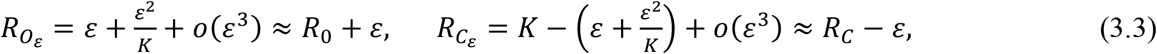

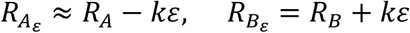, *where k is a positive constant*.

Comparing coordinates of the corresponding equilibria points of system (1.8) with *ε* = 0 and *ε* > 0, we see that the difference between the corresponding coordinates is of the order *ε*, that is, the difference is small for small *ε*.

It follows from formulas (3.2) that non-trivial equilibria *O*_*ε*_ and *C*_*ε*_ exist only if 0 < *ε* < *K*/4 and equilibria *A*_*ε*_ and *B*_*ε*_ exist only if 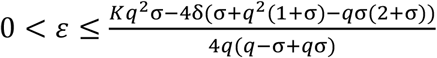, where the equalities correspond to the fold bifurcation in system (1.8).

The system (1.8) depends continuously on the parameter *ε*. Due to this property, the phase structures of hyperbolic equilibrium points of the system with small *ε* > 0 are topologically the same as those for *ε* = 0 . Thus, the points *A,B,C* of system (2.1) and *A*_*ε*_*, B*_*ε*_*, C*_*ε*_ of system (1.8) correspondingly are topologically equivalent for small *ε*, as well as their bifurcation diagrams (up to bifurcations of co-dimension 1). Thus, the point *A*_*ε*_ is a stable topological node, whereas *B*_*ε*_ and *C*_*ε*_ are saddles (see Proposition 2).

Consider now equilibrium points *O* and *O*_*ε*_ in system (1.8). The point *O* is non-hyperbolic in system (2.1), it has a family of homoclinics (elliptic sector) in its non-negative vicinity (see Proposition 4). Including the term *εR* implies “dividing” the point *O* into two equilibrium points, *O* and *O*_*ε*_ in system (1.8). Using standard Trace-Determinant method (see, e.g., [37]) one can show that the point *O*_*ε*_ is a hyperbolic unstable node and the point *O* is a stable (non-hyperbolic) node (see Proposition 6).

As a result, the homoclinics of system (2.1) in the vicinity of *O* are transformed to heteroclinics that connect the points *O* and *O*_*ε*_ (see Fig.6).

So, the following statement holds:

#### Proposition 6.

*Let q* > σ. *Then system (1.8) with ε* > 0 *for all non-negative values of the system parameters has two hyperbolic points, attractive node O and repelling node O*_*ε*_ *, which is placed in ε* − *vicinity of O in the axis R*.

The following Figure 6 shows the difference in the structures of equilibria of systems (2.1) and (1.8).

a) System (2.1); the vicinity of *O*(0,0) consists of an elliptic sector together with stable and unstable node sectors.

b) System (1.8) with *ε* > 0; the system has attractive node *O* and repelling node *O*_*ε*_ that are heteroclinically connected.

**Fig. 6.**
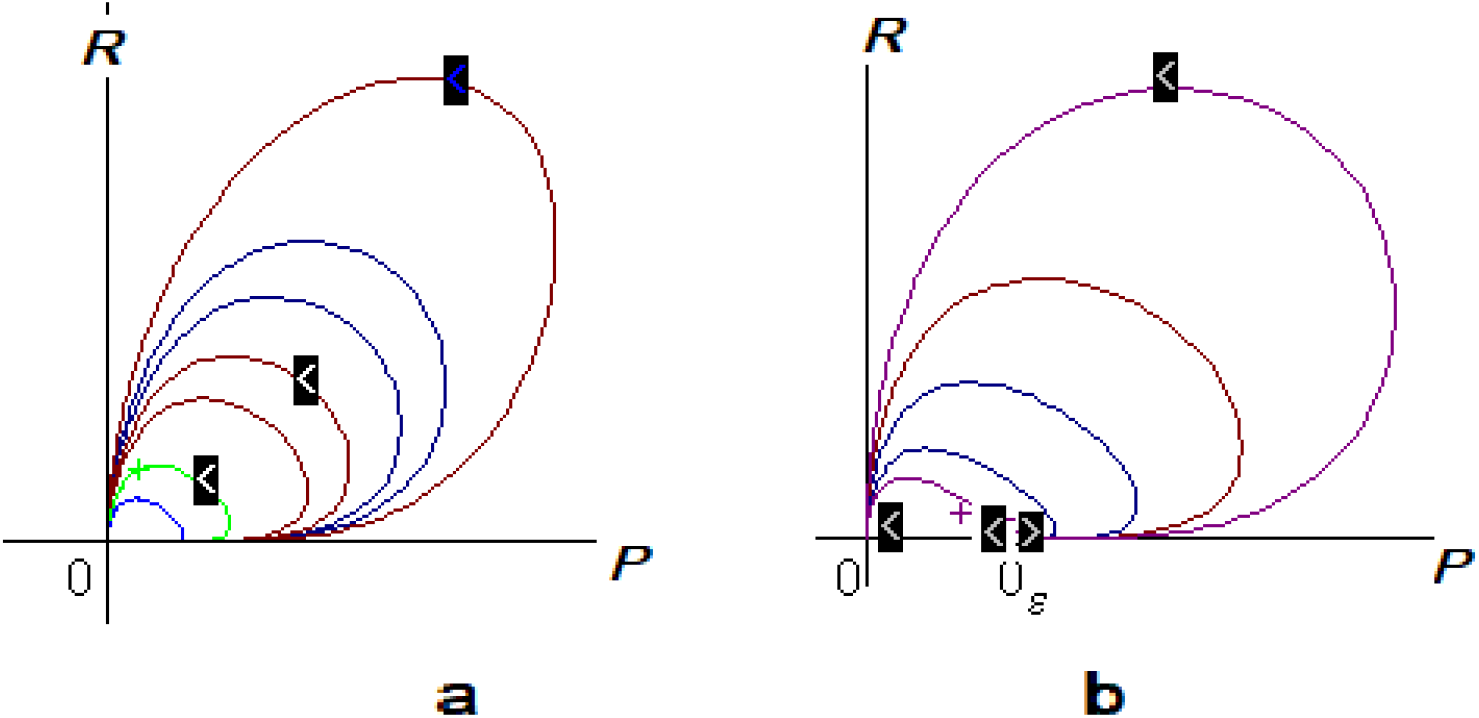
Structure of equilibria of systems (2.1) and (1.8) for *K* = 10, *γ* = 1, *q* = 5.45, *σ* = 2, *δ* = 3;

## Discussion

Here we explore a ratio-dependent model [6] of the simple replicator-parasite system where the parasite emerges as a defective form of the original replicator [4]. Unlike standard two-dimensional prey-predator (host-parasite) models, the ratio-dependent model applies within a broad range of both the replicator and parasite abundances including the regime with very few replicators or parasites. We have shown previously that, within the framework of prey-dependent models, replicators and genetic parasites can coexist in a stable state, but only in a bounded area of the model parameters [36]. The present analysis of the ratio-dependent model shows that the replicators and genetic parasites can either coexist in a stable equilibrium or both go extinct such that the area of coexistence is unbounded. The parasite-free state is unstable in a broad area of the space of the model parameters which include replicator carrying capacity *K*, parasite damage δ, mortality σ, parasite advantage *q* and time scale parameter γ.

More specifically, the two populations coexist in a stable equilibrium state with non-zero numbers of both replicators and parasites if the carrying capacity of the system is large, exceeding the threshold level that is determined by a combination of the other parameters of the system (see Proposition 2), and the parasite advantage parameter *q* is greater than the parasite mortality parameter *σ*. The parametric boundary *q=* σ is of critical importance. The non-trivial positive stable equilibrium *A* appears in the system at *q=* σ and exists for *q>*σ. Simultaneously, for *q*≥ σ, another equilibrium *B* appears and exists in the model; *B* is a saddle whose separatrices are boundaries between the domains of attraction of the non-trivial equilibria *A* (coexistence) and the trivial equilibria *O*(0,0) (extinction). The parasite-free equilibrium *C*(*K,*0) becomes a saddle for *q>*σ. Thus, the parasite-free state of the system is unstable because appearance of even a small number of parasites results in either coexistence or in extinction of both populations. Therefore, the dynamics of the model supports the hypothesis of the inevitable emergence and persistence of genetic parasites due to the evolutionary instability of parasite-free states [4] for an unbounded domain of the parameter space. As the carrying capacity *K* of the system or the parasite mortality rate σ decrease or the parasite damage δ increases, the dynamics of the system is characterized by a competition between the regimes of coexistence and extinction. Specifically, with these changes in the parameter values, the basin of attraction of the non-trivial equilibrium *A* (which can become a spiral) decreases, whereas the basin of attraction of the trivial equilibrium *O* increases. In other words, under these conditions, the extinction of both the replicator and the parasite becomes an increasingly likely evolutionary outcome.

A notable property of the model is that the system presents a dynamical regime of deterministic extinction which corresponds to the existence of a family of homoclinics to the origin, the so-called elliptic sector (Fig. 3). Such a sector can exist only if the origin is a singular equilibrium. The behavior of the system in this regime appears counter-intuitive: both populations start growing from very small initial numbers up to high values, but then, both populations decrease in size and go extinct without any visible reasons, solely due to the internal system dynamics. Notice that this regime disappears if one considers an additional mortality of replicators (system (1.8)). In this case, the singular origin is “divided” onto two simple equilibria resulting more standard dynamics.

In summary, we show here that, within the framework of a ratio-dependent model of replicator-parasite coevolution, the parasite-free state is unstable because appearance of even a very small number of parasites results either in stable coexistence or in the collapse of the entire system. As shown in many previous studies (see [9, 10] for review), ratio-dependent models are better suited for the exploration of host-parasite (prey-predator) systems where the host and parasite population sizes are of different orders of magnitude. In the toy models analyzed here and in our previous work in this area [4, 36], the small parasite population (that is, the high host/parasite ratio) corresponds to the *de novo* emergence of the parasite. However, more generally, this situation would correspond to the arrival of the parasite from the outside, that is, effectively, to any case of infection of a host population. Together with the previous analyses indicating that genetic parasites could not be purged from prokaryote populations due to the high rate of horizontal gene transfer that is essential to the survival of these populations [39, 40], the present results support the concept of genetic parasites as an intrinsic feature of life.

## Author contributions

G.P.K. and F.B. initiated the study; F.B. and G.P.K. developed and analyzed the model; G.P.K. and E.V.K analyzed the results and wrote the manuscript that was read, edited and approved by all authors.

## Acknowledgements

The authors’ research is supported by the Intramural Research program of the National Institutes of Health of the USA (National Library of Medicine).

## Appendix 1. Proof of Proposition 1

To prove the Proposition, we consider the system on the Poincare` sphere (Andronov et al., 1973) and show that there are no stable points in its equators.

We apply the changes of variables in the system (1.8)

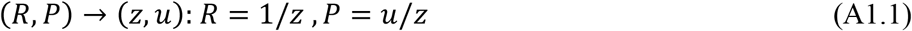

that transforms the “point” *R* = ∞ to axis *z* =0, and

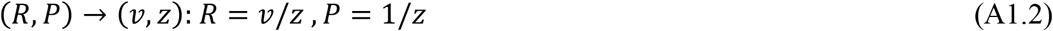

that transforms the “point” *P* = ∞ to axis *z* =0.

Due to transformation (A1.1) supplemented by changing of independent variable *t* by *δ*

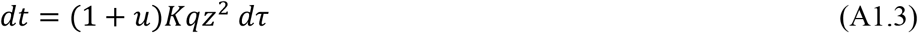

system (1.8) is transforming to

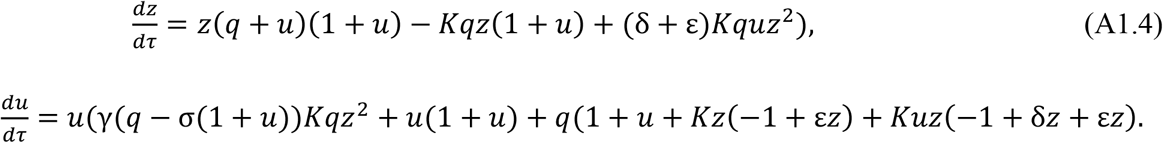

System (A1.4) has a single non-negative equilibrium for *z* = 0, it is the point (*z*, *u*) = (0,0). Simple calculations show that the point (0,0) has eigenvalues *l*_1_ = *q*, *l*_2_ = *q*. It means that the point (0,0) is a *repelling* node.

Due to transformation (A1.2) supplemented by changing of independent variable

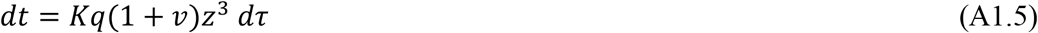

system (1.8) is transforming to the system

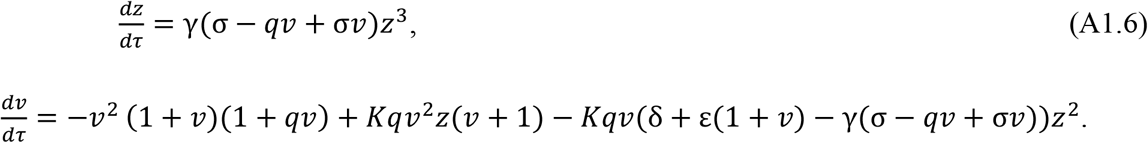

The point (*z*, *v*) = (0,0) is non-hyperbolic, however it has a *saddle sector* in its positive vicinity.

So, we can conclude that all equilibria in equators of Poincare sphere are repelling. The axes *R* = 0 and *P* = 0 are phase curves of the model, so any phase curve starting from a positive point (*R*_0_, *P*_0_) cannot leave the positive quadrant. *The statement is proven.*

## Appendix 2. Proof of Proposition 2

Statements 1) and 2) of the proposition were already proven. Let us prove the statements

3) *Equilibria A, B appear/disappear in the positive quadrant on the boundary **Q**: q* = *σ where the point A coincides with C and the point B coincides with O* (*see Fig.2B*).

4) *Positive equilibria A, B appear/merge composing unique point AB (R*_*AB*_, *P*_*AB*_) *with coordinates* 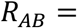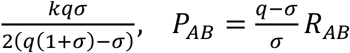 in the boundary

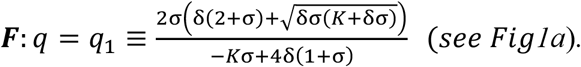

Substituting *q* = *σ* to formulas (2.3), (2.4), one can see that non-hyperbolic point *O* coincides with the point *B* and the point *C* coincides with the point *A*. Both points *O* and *C* exist in the first {*R*, *P*} − quadrant for *q* ≥ *σ* . So, the line *Q*: *q* = *σ* is the boundary where positive equilibria *A* and *B* appear in the phase portrait of the model. Close to this boundary, for *q* > *σ*, the system has four equilibria, *O* and *C*, *A* and *B* in the first quadrant.

Let now *q* = *q*_1_ > σ for fixed σ, δ. It is possible only for *K*σ − 4δ(1 + σ) < 0.

The points *A*(*R*_A_,*P*_A_), *B*(*R*_B_,*P*_B_) merge when 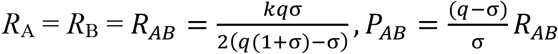.

According to (2.4) it happens with parameter values on the line

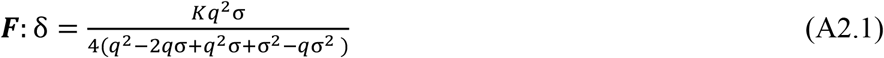

or equivalently on the line:

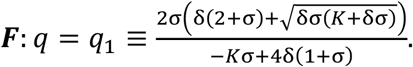

For *q* > *q*_1_ the system has no nontrivial equilibria. Proposition is proven.

## Appendix 3. Proof of Proposition 3

Jacobian *J*(*R*, *P*) of system (2.1) is of the form:

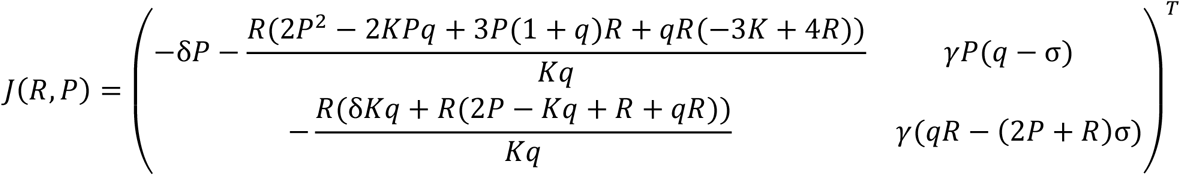

1) Substituting coordinates of the point *C* to *J* we get 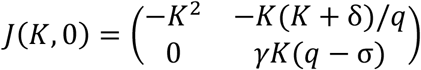

Its eigenvalues are *μ*_1_ = −*K*^2^, *μ*_2_ = *γK*(*q* − σ). So, the point *C* is a saddle for *q* > σ and a stable node for *q* < σ. Eigenvalue *μ*_2_ = 0 for *q* = σ, so the transcritical bifurcation happens on the line *q* = σ; the equilibrium *C* is a saddle–node here.

2) We analyze structures of the equilibria *A* and *B* using standard Trace-Determinant method (see, e.g., [37] or [41]). It is known that an equilibrium is a saddle if the determinant of Jacobian in this point is negative, and “non-saddle” (a node, spiral/center) if it is positive. A “non-saddle” equilibrium is stable if the trace of the Jacobian in this point is negative and unstable if the trace is positive; the case of zero trace can correspond to Andronov-Hopf bifurcation [41].

Consider equilibria points *A* (*R*_A_, *P*_A_)*, B* (*R*_B_, *P*_B_):

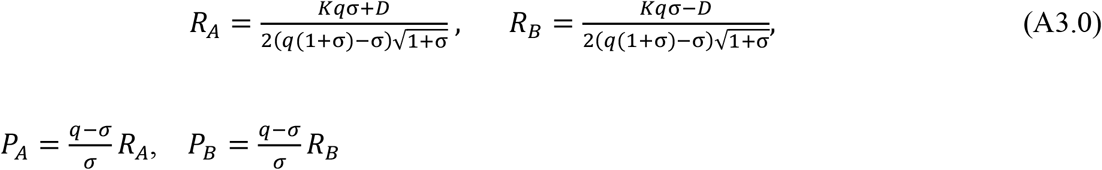

where 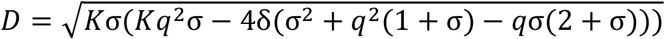.

Substituting *P* = *P*_*A*_ or *P* = *P*_*B*_ to *J*(*R*, *P*) we get the matrix

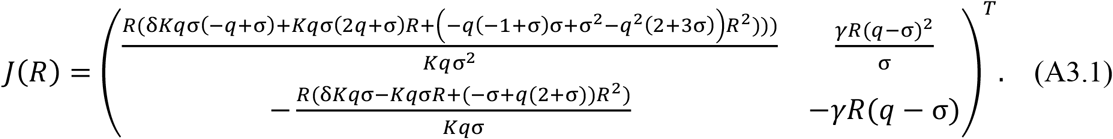

where *R* = *R*_*A*_ or *R* = *R*_*B*_.

Determinant *Det*(*R*) of Jacobian *j*(*R*) is of the form

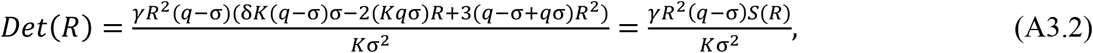

where *S*(*R*) = 2δ*K*σ(*q* − σ) − 3*Kq*σ*R* + 4(*q* − σ + *q*σ)*R*^2^;

the trace of the Jacobian *J*(*R*)

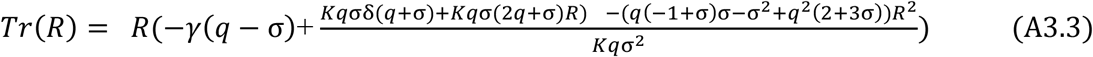

for both *R* = *R*_*A*_ or *R* = *R*_*B*_. Substituting *R* = *R*_*A*_, *R* = *R*_*B*_ to *S*(*R*) one gets

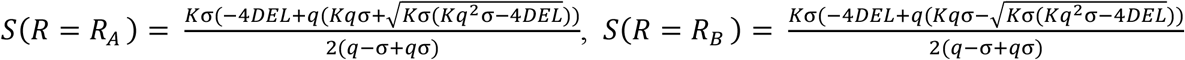

where *DEL* ≡ 4δ)*q* − σ)(*q*(1 + σ) − σ)) ≤ *Kq*^2^σ.

It is easily to verify (e.g., with the help of MATEMATIKA-package) that

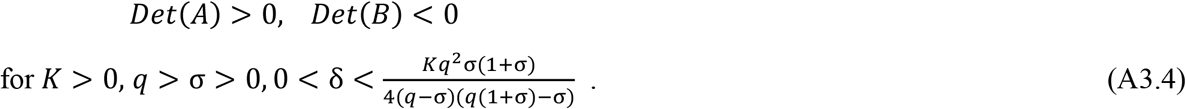

Thus*, the point A*(*R*_*A*_, *P*_*A*_) *is a topological node (node/spiral/center), and B*(*R*_*B*_, *P*_*B*_) *is a saddle in the domain* (A3.4) *of their existence*.

3) Consider now the structure of equilibrium *A.* Let us present the trace of matrix (A3.1) in equilibrium *A* as a sum:

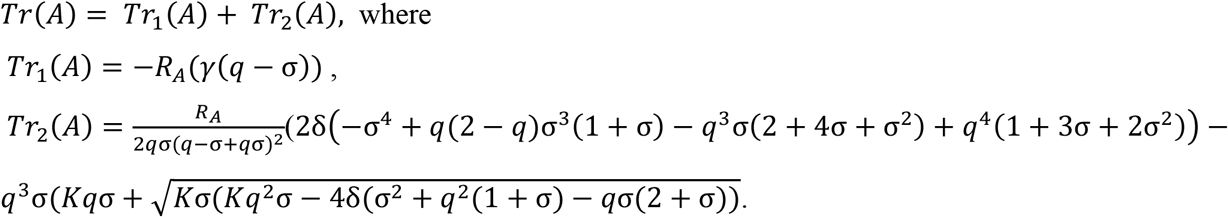

Evidently, *Tr*_1_(*A*) < 0 if *q* > σ, in this case *A* belongs to the first quadrant (see (2.7) or (A3.0)).

It is possible to verify (e.g., using the MATEMATIKA-package) that *Tr*_2_(*R*_*A*_) < 0 for

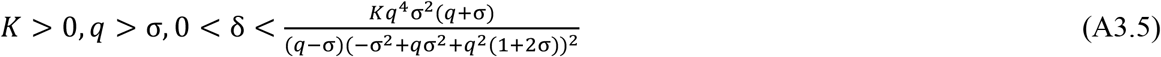

and *Tr*_2_(*A*) > 0 for

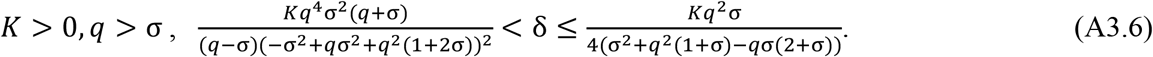

*Tr*(*A*) < 0 in the first case, so *A* is a stable topological node, *Tr*(*A*) can change the sign in the second case (for example, at large *γ*) and for some parameters we can get *Tr*(*A*) ≥ 0.

Consider now the case *Tr*(*A*) = 0. Substituting *R*_*A*_ from (2.8) to (*A*3.3) and solving equation *Tr*(*A*) = 0 with respect to δ we obtain the boundary

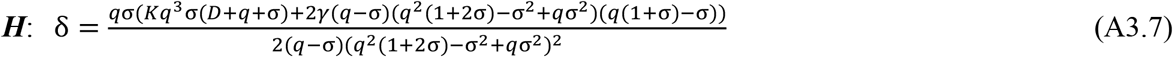

where 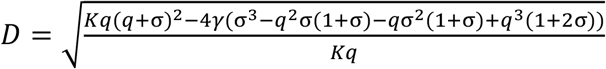

Notice that with parameter values (A3.7) the equilibrium *A* can change stability *via the Andronov-Hopf bifurcation*. It can happen only in Domain II where *K*σ − 4δ(1 + σ) < 0 (see Fig.2).

Numerical analysis of the model at parameter values close to *H* has shown that *subcritical* Hopf bifurcation is realized in the system.

4) Now we apply the method of small parameter to analyze the structures of equilibria *A* and *B* close to boundaries of their existence in the 1^st^ quadrant. Consider firstly crossing the line *q* = σ. Expanding *R*_*A*_, *R*_*B*_ and *Det*(*A*), *Tr*(*A*) in Taylor series with respect to *d* = *q* − σ we get

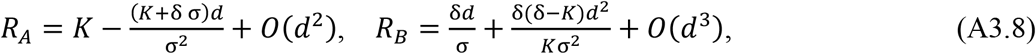

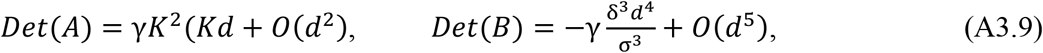

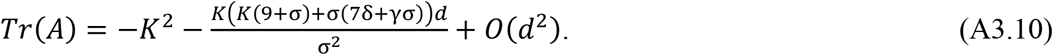

We see that *Det*(*A*) > 0, *Tr*(*A*) < 0 for small positive *d* = *q* − σ. So, the equilibrium *A* is a stable node for small positive *d* and a saddle for *d* < 0.

Thus, the points *A* and *C* exchange stability on the line ***Q***: *q* = *σ*, i.e. the *transcritical* bifurcation happens in the system.

*Remark*. For the point *B* given by (A3.8), *Det*(*B*) < 0 if *d* is small; so, the equilibrium *B* is a saddle as we have already proven (see (A3.4)).

The equilibria *A*, *B* merge on the boundary 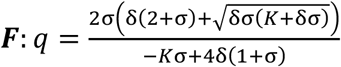 composing the point *AB*(*R_AB,_ P_AB_*) (see Fig. 3). Let us consider the structure of the point *AB*.

From equation (2.8) we get 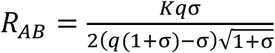. Substituting this value of *R* to (A3.3) we get

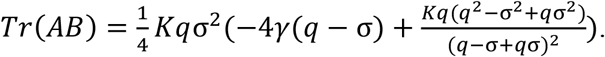

Solving equation *Tr*(*AB*) = 0 we get 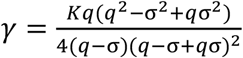

Thus, the system of equations

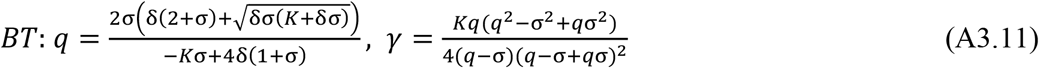

defines the *Bogdanov-Takens (BT-) bifurcation* of co-dimension 2 which is realized in the model.

So, the point *AB* where equilibria *A* and *B* merge can be a saddle-node; the point *AB* is a degenerate saddle for parameter values (A3.11) (see, e.g., [38]).

We also verify numerically that *BT- bifurcation* realizes in the system. We have observed the unstable limit cycle that appears from the “separatrix loop” composed by separatrices of the saddle *B.* At further changing of parameters the cycle “sets” to the stable equilibrium *A* that becomes unstable.

## Appendix 4. Proof of Proposition 4

Due to Propositions 2 and 3 the equilibria of model (2.1) are placed in the first quadrant with all positive parameters *K*, *γ*, *δ*, *ϵ*, *q* ≥ *σ* and only equilibrium points can serve as attracting manifolds. There are two cases of placing of the equilibria for considering parameters (see Fig 1). The system has two equilibria, the *saddle C*(*K*, 0) and *non-hyperbolic* point *O*(0, 0), if *Kσ* − 4*δ*(1 + *σ*) < 0, 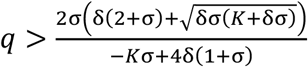 and four points *O*(0,0), *C*(*K*, 0), *A, B* if *Kσ* − 4*δ*(1 + *σ*) > 0 or if

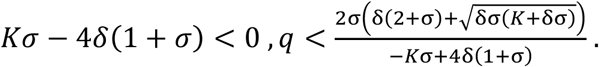

For analysis the structure of *O* we apply the “blowing-up method” that allows to divide non-hyperbolic point to some hyperbolic ones [42, 43]; some results proven in these papers are collected in the following

### Proposition A

1. *a vicinity of non- hyperbolic equilibrium point contains an elliptic sector if one of blown-up systems has neighboring attractive and repelling nodes or if the origin in one of the systems is attractive node and the origin in another system is a repelling node;*
2. *a vicinity of non- hyperbolic equilibrium point contains a parabolic sector if one of blown-up systems has neighboring a node and a saddle or if the origin in one of the systems is a node and the origin in another system is a saddle;*
3. *a vicinity of non- hyperbolic equilibrium point contains a hyperbolic sector if one of blown-up systems has neighboring saddles or if the origins in both systems are saddles.*

These statements are used for proving the Propositions 4 and 6.

A behavior of system (2.1) in a vicinity of *O*(0,0) is defined by the system

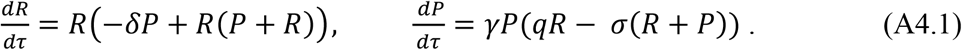

For analysis of the structure of equilibrium *O*(0,0) of system (A4.1), we apply two blowing-up changes of variables:

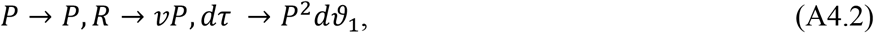

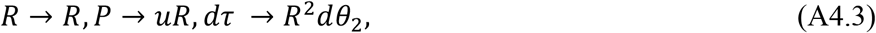

that transform the point *O* to the axes *P* = 0 and *R* = 0 correspondingly.

System (A4.1) is transformed by changing (A4.2) to the system:

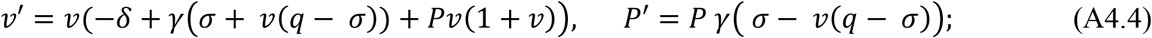

changing (A4.3) transforms (A4.1) to the system:

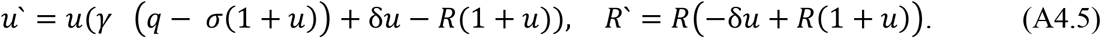

Let us consider system (A4.4). The system has equilibria *o*_1_(*P* = 0, *g* = 0) and 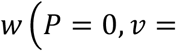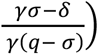. The point *w* is placed in the positive quadrant if *q* > *σ* and *γσ* > *δ*.

Equilibrium *o*_1_ has eigenvalues *l*_1_(*t*_1_) = −*γσ*, *l*_2_(*o*_1_) = −*δ* + *γσ*, so *o* _1_ is an attractive node if *γσ* < *δ* and is a saddle if *γσ* > *δ*. Equilibrium *w* has eigenvalues *l*_1_(*α*) = *δ* − *γσ*, *l*_2_(*α*) = −*δ*_;_ so *w* is an attractive node in the first quadrant for *q* > *σ* and *γσ* > *δ*; *w* is out the first qudrant if *γσ* < *δ*.

Consider now system (A4.5). This system has equilibrium *o*_2_(*u* = 0, *R* = 0), which is a repelling node. (A4.5) has also equilibrium point 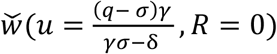 that belongs to the first quadrant for *q* > *σ* and 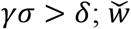 is out the first qudrant if *γσ* < *δ*. The equilibria *w* and 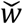 has the same structures. So, we need to analyze two cases *γσ* < *δ* and *γσ* > *δ*. In both cases we apply *Proposition A* for proving existence of an elliptic sector in a vicinity of the point *O* of system (A4.1)

In the first case the blown-up systems (A4.4) and (A4.5) have, correspondingly, attracting and repelling node in their origins. Thus, system (A4.1) has an elliptic sector due to Proposition A, 1) with asymptotic boundaries *R* = 0 and *P* = 0. In the second case systems (A4.5) has neighboring equilibria, attracting node 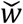 and repelling node *o*_2_. Thus, system (A4.1) has an elliptic sector due to Proposition A, 1) with asymptotic boundaries *R* = 0 and 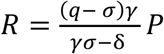 (see Fig.3).

Proposition 4 is proven.

## Appendix 5. Proof of Proposition 6

Consider system (1.8) with *ε* > 0. The system has at most five equilibria in the first quadrant whose coordinates given in (3.2), (3.3). A new equilibrium *O*_*ε*_ has coordinates 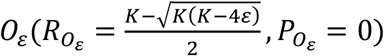 for 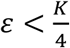, and 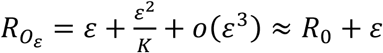.

Equilibrium point *O(0,0)* of system (1.8) is *non-hyperbolic* for all *ε* ≥0 because eigenvalues of *O(0,0)* are equal to zero.

Let us consider the structure of the point *O(0,0)* in system (1.8) with *ε* > 0 by blow-up method. The following system keeps the main terms of system (1.8) in a vicinity of *O:*

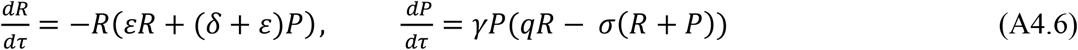

Changing of variables (A4.2) and (A4.3) transform this system correspondingly to equations

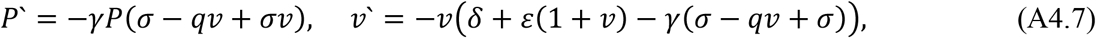

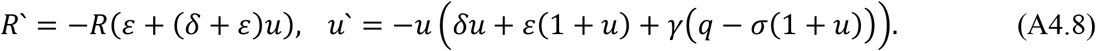

It is possible to show that non-negative equilibria of (A4.7) in the axis *P* = 0 are consequent stable node and a saddle. Thus (see Proposition A.2) system (A4.6) has stable parabolic sector in a vicinity of equilibrium *O*.

System (1.8) has also equilibrium 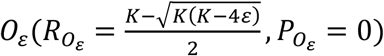 (see (A3.2)). Using the Trace-Determinant method one can easily show that equilibrium *O*_*ε*_ is a repelling node (see Fig.3 b).

**Proposition is proven.**

## Notes

### Competing Interest Statement

The authors have declared no competing interest.

